# Modeling the emergence of antibiotic resistance in the environment: an analytical solution for the minimum selection concentration

**DOI:** 10.1101/176289

**Authors:** Ben K. Greenfield, Shanna Shaked, Carl F. Marrs, Patrick Nelson, Ian Raxter, Chuanwu Xi, Thomas E. McKone, Olivier Jolliet

**Affiliations:** Department of Environmental Health Sciences, University of California - Berkeley, CA, USA; Department of Epidemiology, University of Michigan, Ann Arbor, MI, USA; Department of Environmental Health Sciences, University of Michigan, Ann Arbor, MI, USA; Department of Mathematicsd, University of Michigan, Ann Arbor, MI, USA

## Abstract

Environmental antibiotic risk management requires an understanding of how subinhibitory antibiotic concentrations contribute to the spread of resistance. We develop a simple model of competition between sensitive and resistant bacterial strains to predict the minimum selection concentration (MSC), the lowest level of antibiotic at which resistant bacteria are selected. We present an analytical solution for the MSC based on the routinely measured minimum inhibitory concentration (MIC) and the selection coefficient (sc) that expresses fitness differences between strains. We calibrated the model by optimizing the shape of the bacterial growth dose–response curve to antibiotic or metal exposure (the Hill coefficient, κ) to fit previously published experimental growth rate difference data. The model fit varied among nine compound-taxa combinations examined, but predicted the experimentally observed MSC/MIC ratio well (R^2^ ≥ 0.95). The shape of the antibiotic response curve varied among compounds (0.7 ≤ κ ≤ 10.5), with the steepest curve for the aminoglycosides streptomycin and kanamycin. The model was sensitive to this antibiotic response curve shape and to the sc, indicating the importance of fitness differences between strains for determining the MSC. The MSC can be more than one order of magnitude lower than the MIC, typically by a factor sc^κ^. This study provides an initial quantitative depiction and a framework for a research agenda to examine the growing evidence of selection for resistant bacteria communities at low environmental antibiotic concentrations.

## Introduction

Effective management of antibiotic risks in the environment requires an understanding of the factors responsible for the emergence, transmission, and maintenance of antibiotic resistance (1). It is particularly important to address the question of when resistant bacteria predominate as a result of environmental antibiotic pollution (1–5). For example, insights are also needed into the extent to which antibiotics in aquatic environments contribute to the spread of resistance, and to the long-term prevalence of resistant infections in humans (2, 5).

The mutant selection window (MSW) paradigm states that resistant mutants may develop between the lowest boundary concentration of selection for resistance, and the upper boundary concentration of growth inhibition of the most resistant potential mutant (the mutant prevention concentration, MPC) (6, 7). The paradigm further indicates that the lower boundary concentration of the MSW is the minimum concentration that inhibits colony formation (MIC, ng ml^-1^), and the MIC has been useful to evaluate hazard of selection for resistance in natural aquatic environments (4, 8). Considerable research *in vitro* and *in vivo* has demonstrated that resistant mutants develop between the MIC and the MPC (7, 9, 10), but many laboratory and theoretical studies indicate that resistant mutants can also be preferentially selected above the minimum selective concentration (MSC, ng ml^-1^), defined as the lowest concentration at which a resistant strain outcompetes and displaces sensitive isolates (1, 11–18). Because the MSC can be lower than the MIC, and to minimize the hazard of resistance occurring in the natural environment (e.g., aquatic systems), further characterization and understanding of the MIC versus MSC relationship would be beneficial (8).

Laboratory experiments (11–13, 16, 18) have elegantly demonstrated MSCs ranging from 1/4 to below 1/200 of the MIC for antibiotics of several classes (e.g., macrolide, aminoglycoside, fluoroquinolone, and antifolate) and for two metals in *Escherichia coli* (*E. coli*) or *Salmonella enterica* serovar Typhimurium LT2 (*S.* Typhimurium). This finding may help explain the high levels of resistance found in the environment, particularly at subinhibitory antibiotic concentrations (2, 5, 11). These studies further indicate that the fitness cost of the resistance-conferring mutations is more important than differences in MIC between strains for discerning how much below the MIC the resistant bacteria will predominate (12). However, a mathematical description of the competition between strains would aid in understanding strain- and antibiotic-specific results and generalizing to a wider range of situations.

Mechanistic mathematical models, including experimentally validated pharmacodynamic/ pharmacokinetic models, describe antibiotic effects better than simple MIC measurements (19–23). For example, the shape of the antibiotic dose–response curve is very important to the microbiological efficacy of antibiotic treatment regimens at high (treatment) levels (19). The implications of this understanding of dose–response curve shape for low (subinhibitory) antibiotic levels and for calculation of the MSC, while relevant for selection of resistance, have not been considered in as much depth. To complement the recent empirical research (1, 11–13, 16, 18), there remains a need for a quantitative model describing the MSC, i.e., the minimum environmental antibiotic concentration that allows resistant bacterial strains to dominate. Such a model can generate testable predictions, identify the factors that determine water or soil antibiotic concentrations that select for resistance, and be incorporated into risk assessments of antibiotic resistance development (1).

An analytical solution for the MSC has two potential uses. First, model sensitivity analysis and examination of parameter structure may provide insight on the relationship between commonly considered bacterial growth and antibiotic dose–response parameters and the MSC itself. Second, current methodology to accurately measure the MSC requires direct measurement of competition between bacterial strains and specialized methods such as fluorescent cell tagging and flow cytometry (12, 16). An analytical solution provides a potential alternative to these methods, instead estimating the MSC based on bacterial growth rate and antibiotic dose–response parameters that are routinely obtained within microbiology laboratories. To that end this paper addresses three questions: 1. How do we quantitatively define the minimum selective concentration (MSC) via a parsimonious mathematical model in combination with readily available measurements? 2. How well does such a mathematical model of MSC fit to published empirical data? 3. What model parameters, representing biological characteristics, are most important to describe the MSC?

The model we propose in this paper describes the MSC based on the competition between a wild-type and a resistant strain of bacteria, and the key factors that favor the growth of resistant strains at subinhibitory antibiotic concentrations. The model focuses on conspecific gram negative bacteria (GNB), and is calibrated to the experimental results of Gullberg et al. (12, 16) for *E. coli* and *S.* Typhimurium. The model illustrates the shape of the antibiotic dose–response curve as a measurable and influential driver on the ratio of the MSC and MIC, and presents a hypothesized dose–response relationship for use in risk assessment of resistance development in environmental settings. Finally, we discuss the implications of the MSC results for increased risk of antibiotic resistance selection at antibiotic concentrations observed in antibiotic-contaminated waste streams and natural waters.

## Theory

We develop a simple analytical expression of the ratio between the MSC and the MIC for a sensitive strain (i.e., MSC/MIC), which mathematically describes the factors that determine risks of subclinical antibiotic concentrations (11, 12). The model is based on the competition between two bacterial strains: a wild type sensitive strain, and mutant strain that is more resistant.

### Model derivation for net growth rate

At a given antibiotic concentration *a* [ng ml^-1^], the net growth rate (N(*a*), [h^-1^]) for each strain is given by

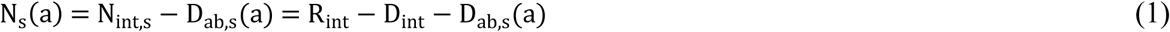

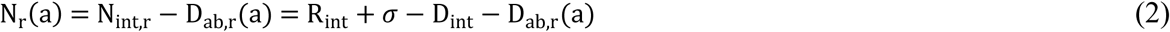

where subscript s = sensitive bacteria, subscript r = resistant bacteria, N_int,s_ = (R_int_ – D_int_) = intrinsic net growth rate in the absence of antibiotic [h^-1^],, calculated as the difference between the intrinsic growth rate (Rint =[ h^-1^]) and the intrinsic loss due to mortality (or, in continuous cultures, dilution) (Dint, [h^-1^]), D_ab_(*a*) = loss in net growth [h^-1^] due to a given antibiotic concentration, *a*, and σ = absolute selection coefficient [h^-1^].

The absolute selection coefficient (σ) represents the loss in fitness of resistance-conferring genes as the absolute difference in net growth rate between bacteria strains (e.g., sensitive vs. resistant) in the absence of antibiotics (i.e., N_int,r_ = N_int,s_ + σ). The absolute selection coefficient (σ) is directly related to the fitness cost (see supplemental material about the exact definition of fitness cost and its relation to the absolute selection coefficient). Accurate measurement of the absolute selection coefficient (σ) can be difficult, employing competition experiments with labeled strains and flow cytometry (12, 16, 24). Resistance-conferring mutations exhibit highly variable selection coefficients in comparison to sensitive strains (24–27), with compensatory mutations often reducing or reversing the fitness cost of resistance mechanisms (26, 28, 29). For the purposes of this model, we run simulations on the assumption that resistance-conferring mutations engender a loss in fitness, resulting in lower growth rates relative to less resistant strains, i.e., σ < 0 in Eq. 2. This assumption is supported in that the majority of single mutational events entail a loss in fitness (24).

The loss in net growth due to antibiotics can be described by a generalized Hill Equation (19, 21, 30–32):

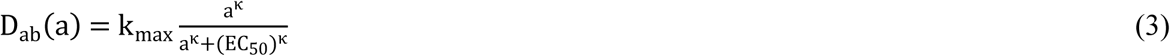

in which k_max_ [h^-1^] is the maximum death rate due to antibiotic, EC_50_ [ng ml^-1^] is the antibiotic concentration that achieves half of this maximum rate, and will thus increase with increased resistance, and κ is the Hill coefficient, which gives an indication of how steeply D_ab_ increases near the MIC (31). For κ = 1 in the range of antibiotic concentrations below the MIC, the death rate increases roughly linearly. For a given strain, antibiotics with a high κ value (> 1) will have lower efficacy at sub-therapeutic levels, but higher efficacy at therapeutic levels above the MIC. The opposite relation is true for antibiotics with low κ values (19) as illustrated in Fig. 1A.

**Fig. 1.**
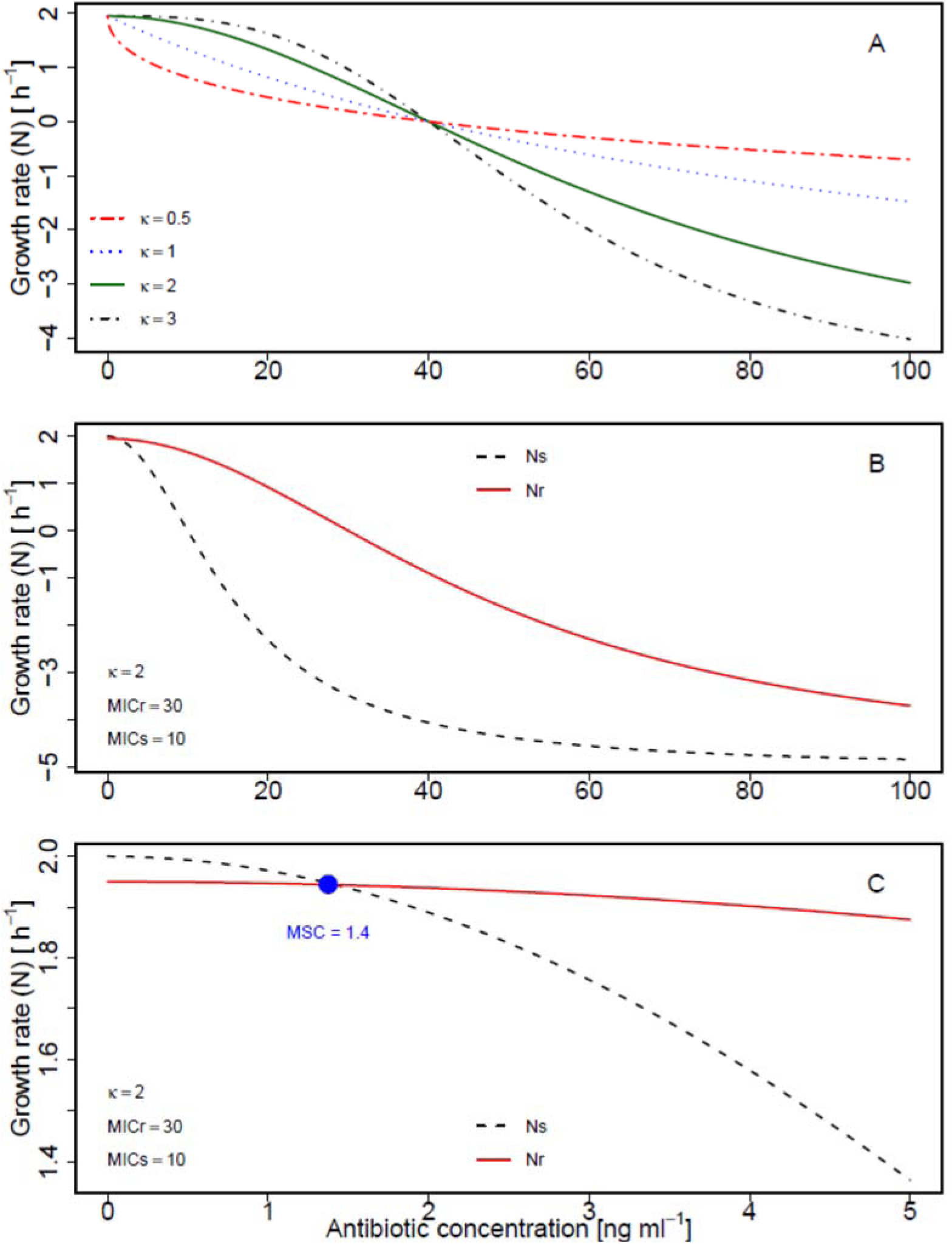
Growth rate versus antibiotic concentration. (A) Single strain (MIC_r_ = 40) with different kappa (κ) values. (B, C) Sensitive and resistant strains (MIC_s_ = 10, MIC_r_ = 30, κ = 2). (C) enlarged view around the MSC (•), the concentration where growth curves cross (ΔN = 0). Other parameter values: selection coefficient = 0.05, N_int,s_ = 2, N_min_ = −5.

To determine k_max_ from growth and death rates, we note that k_max_ should correspond to the difference between the maximum possible net growth rate (not limited by resource availability or antibiotics; i.e., N_int_), and the minimum possible growth rate, after accounting for the growth limiting activity of antibiotic (N_min_):

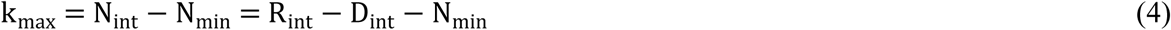

Generally, N_min_ < 0, indicating population decline at maximum antibiotic exposure level. The EC_50_ can be directly related to the MIC value [ng ml^-1^]; as a result, the following formulation of D_ab_ applies for our formalism (full derivation in the supplemental material):

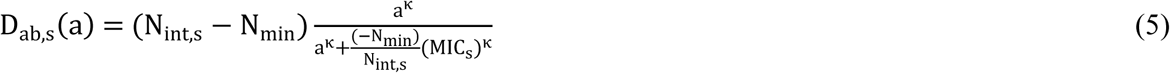

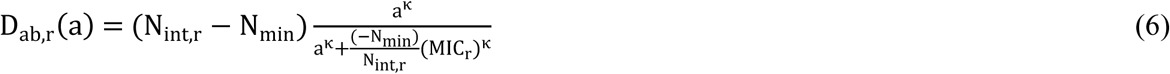

Equations 5 and 6 assume identical κ and N_min_ for sensitive versus resistant strains, which may not be accurate. Later in the text, we revisit the impact of this assumption for estimation of the MSC.

### Difference in net growth rate and derivation of MSC as function of MIC

Competition between different bacterial strains is expressed by the difference in net growth rates. According to the conceptual model described by Andersson and Hughes (11) and Gullberg et al. (12), N_s_ > N_r_ at low antibiotic concentrations, but the greater sensitivity causes more antibiotic-dependent growth inhibition for the sensitive strain. As a result, at high antibiotic concentrations, N_r_ > N_s_, and the MSC is the point of intersection of the two growth curves (N_s_ = N_r_) for which the difference in net growth rate is zero (Fig. 1B-C).

Analytically, this difference in net growth rates between the resistant and the sensitive strain (ΔN(a) [h^-1^]) is determined by subtracting Eq. 1 from Eq. 2, giving:

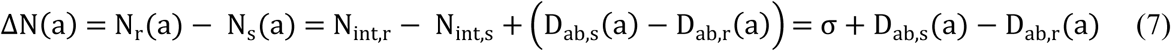

Thus, the MSC is the antibiotic concentration (i.e., a = MSC) at which the two net growth rates are equal and the difference (Eq. 7) is zero:

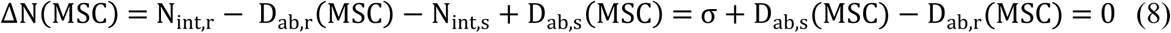

This is the concentration at which the additional loss in net growth due to antibiotic in the sensitive strain compared to the resistant (D_ab,s_(MSC) - D_ab,r_(MSC)) compensates for the effect of fitness loss (σ).

To derive the ratio of MSC/MIC we employ a dimensionless relative selection coefficient (sc [unitless]), obtained by reversing the sign of the reported absolute selection coefficient (σ) (12, 16), and then dividing by the net growth rate of the sensitive strain (full derivation in the supplemental material):

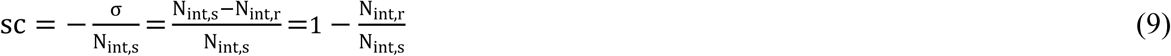

Based on the above equations, and further assuming that κ and N_min_ are the same for sensitive and resistant strains, the following analytical solution for MSC/MIC_s_ is obtained (full derivation in the supplemental material):

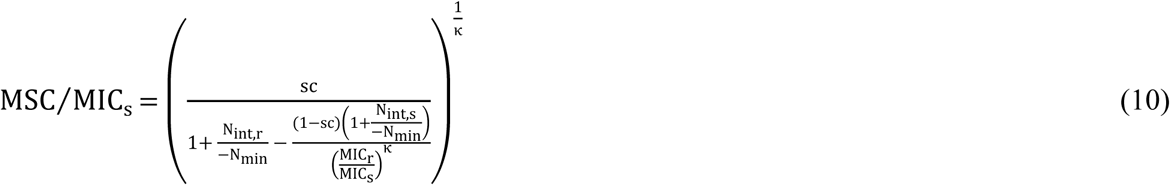

In the case of a large difference in resistant versus sensitive MIC, the right-hand term in the denominator approaches zero, and the equation simplifies to:

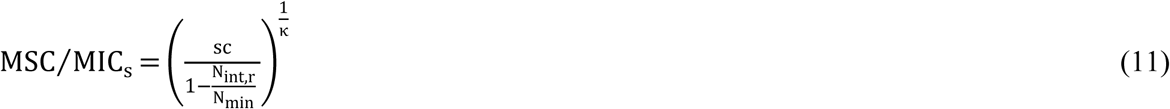

This simplification does not apply to small increases in MIC, such as the *ΔmarR* and *ΔacrR* mutants which double the MIC for ciprofloxacin (12). Eq. 11 becomes appropriate once MIC_r_ > 5 x MIC_s_, at which point results from Eqs. 10 and 11 become approximately equal (supplemental material Fig. S1).

Eq. 10 is, in essence, a mathematical hypothesis regarding the relationship between the antibiotic dose–response and the resulting MSC. The form of Eq. 10 aids in determining which aspects of the growth rate are likely to be most important for competition at low antibiotic doses. Eqs. 10 and 11 also represent a potential alternative to direct measurement of MSC. κ, N_int,s_, and N_min_ could be measured in the laboratory (19, 33 provide examples) and other parameters obtained from literature. Eq. 4 would be fit to experimental measures of the sensitive strain’s antibiotic versus growth dose–response to obtain κ and k_max_, and N_min_ would then be based on the strain intrinsic growth rate (N_int_) minus k_max_ (Eq. 5).

As mentioned above, Eq. 10 rests on the assumption of identical κ and N_min_ for sensitive versus resistant strains. An analytical solution analogous to Eq. 10 could not be obtained assuming separate κ and N_min_ (i.e., κs, κr, N_min,s_, N_min,r_). In the Results section below and supplemental material, we employ a Monte Carlo Simulation sensitivity analysis to critically evaluate this assumption of identical κ and N_min_.

### Model evaluation against experimental results

The analytical solution was evaluated by comparison to the experimental results of Gullberg et al. (12, 16). This evaluation was performed to determine whether the model fit to actual competition data was reasonable, and to identify representative parameter sets for N_min_ and κ, parameters, given a set of published values of MIC_s_, MIC_r_, σ and N_int,s (12, 16)_. The latter four parameters have been characterized for a wide range of strains, conditions, and resistance mechanisms (4, 25, 26, 28, 34–36).

Model fitting was achieved by fitting ΔN values calculated from Eq. 7 (based on Eqs. 1, 2, and 4 - 6) to the ΔN values observed in Gullberg et al. (12, 16). The function NonLinearModel.fit in MATLAB (Statistics Toolbox, R2013a, MathWorks, Natick, MA, USA) was used to estimate N_min_ and κ. Fitting was performed separately for seven individual antibiotic-bacteria combinations across the published range of experimental concentrations, as well as for arsenite- and copper-exposed *E. coli* (12, 16). These metals were included based on co-resistance and cross-resistance with antibiotics, as well as similar mechanisms of genetic transmission among bacteria (2, 5, 37, 38). For each compound, resistance was compared between a sensitive (wild-type) strain and one to four resistant strains in *S.* Typhimurium or *E. coli*. From the published experiments (12, 16), only the chromosomal mechanism of trimethoprim resistance was excluded because it exhibited an average selection coefficient σ > 0, indicating no selective disadvantage of resistance (12).

To evaluate Eq. 7 and the underlying model assumptions, model-predicted vs. observed ΔN were compared. To evaluate robustness to individual observations, cross-validation (CV) was also employed. For CV, the optimization was performed with each single data point removed in series, and the average and range of N_min_ and κ results were examined, as well as the calculated vs. observed ΔN for the out-of-sample observations. The PRESS statistic (predictive residual sum of squares) was calculated, and PRESS/SSY and PRESS/SSE were examined to indicate model prediction error and robustness to individual observations, respectively (39). All analyses were performed on the experimental average results for each strain and antibiotic concentration examined, reported by Gullberg et al. (12, 16). Analyses were also performed on the raw data for each experimental observation (12, 16 supplemental material), in order to consider the impact of experimental variation on results.

## Results

Figure 1 depicts the change in growth rate versus antibiotic concentration (Eq. 7), the crossover point between growth rate of sensitive and resistant strains (i.e., the MSC, Eq. 8), and an analytical solution for the MSC/MIC ratio (Eqs. 10, 11). The MSC can be observed as the antibiotic concentration at which N_s_ (Eq. 1) and N_r_ (Eq. 2) cross, indicating identical growth of sensitive and resistant strains (Fig. 1B-C, Eq. 8). The MSC/MIC ratio (12) is of interest because it indicates how much lower the MSC is relative to the MIC; this enables estimation of the environmental antibiotic concentration at which resistance selection could occur among competing bacteria populations (1, 15). The MSC can be estimated employing this ratio, in combination with routinely available MIC data (4, 8, 34, and the EUCAST database: http://www.srga.org/eucastwt/wt_eucast.htm).

We first evaluate the model by examining a key assumption and then comparing predicted growth rates and MSC/MIC ratios to published data. We then examine model behavior and implications for MSC/MIC ratio prediction. Finally, we perform a sensitivity analysis to identify the most important parameters for predicting this ratio.

### Model evaluation

#### Effect of varying κ and N_min_ for sensitive versus resistant strains

The analytical solution for Eq. 10 requires identical κ and N_min_ for sensitive versus resistant strains. We evaluated the impact of this assumption on model predictions by determining which strain-specific parameter values (i.e., κs, κr, N_min,s_, or N_min,r_) were most important for predicting MSC. To achieve this, we performed a Monte Carlo Simulation sensitivity analysis, detailed in the text and Table S1 (supplemental material). In two simulations, the predicted value of MSC was obtained in Eq. 10, assuming separate κs, κr, N_min,s_, and N_min,r_ in Eqs. 5 and 6. To be robust to MIC ratio variations, the first simulation had MIC_r_ = 1.5 x MIC_s_ whereas the second had MIC_r_ = 10 x MIC_s_. In both simulations, MSC was highly sensitive to κs, (Spearman rank correlation coefficient, ρ > 0.8) but was insensitive to N_min, s_ or N_min, r_ (|Spearman ρ| ≤ 0.11). This much stronger influence of κ than N_min_ is expected based on the fact that κ is an exponential term (Eqs. 10, 11). MSC was also more sensitive to κs than κr, and |Spearman ρ| between κs and MSC (ρ = 0.83) was more than twice |ρ| between κr and MSC (ρ = −0.39). When MIC_r_ = 10 x MIC_s_, almost all variation in MSC was explained by κs (ρ = 0.97), with ρ = −0.09 for κr. These results indicate that MSC will strongly depend on κs, the shape of the antibiotic dose–response for the sensitive strain. As a result, for indirect estimation of MSC using Eq. 10, κs should be well characterized experimentally.

#### Model performance for predicting difference in net growth rates

Figs. 2 and 3 display the difference in net growth rates for sensitive versus resistant strains (ΔN) for previously published experimental data in comparison to the model (Eq. 7). Fig. 2 illustrates how ΔN increases with increasing antibiotic concentration for the ciprofloxacin experiments in Gullberg et al. (12), with variability due to examination of four bacterial strains. Fig. 3 directly compares the experimentally observed versus model predicted ΔN for all antibiotics and metals examined. The model predicted results overlapped with the range of experimental observations for most conditions. Much of the variability was attributable to experimental variation rather than model error, as evident in the horizontal spread of points for each compound. However, the model underpredicted experimental results for the aminoglycosides, KAN and STR (open circles in Fig. 3) when ΔN was < 0.05. Consequently, linear regression indicated ΔN_modeled_ = 0.93(ΔN_observed_) – 0.002, a slight underprediction. Examining results for individual compounds, model performance (R^2^, Q^2^, and PRESS/SSY) was generally similar when either one parameter (κ) or two parameters (κ, N_min_) were fitted, and for either the raw or averaged experimental data (supplemental material Table S2). For CIP, ERY, KAN, and STR, the model fit was insensitive to N_min_, exhibiting a wide range of possible values, and a limited impact on model fit. Therefore, N_min_ was fixed at a representative literature value of N_min_ = −2 (19, 33, 40), and κ was fitted to experimental observations. The fitted model was generally consistent with raw observations (R^2^ > 0.8) and the model exhibited high predictive value in cross validation (Q^2^ > 0.8) for TET, TMP, ERY, and As in *E. coli*, and for TET in Salmonella (Table 1, supplemental material Fig. S2). Model fit was moderate for CIP (R^2^ = 0.78, Fig. 2), Cu (R^2^ = 0.73), and STR (R^2^ = 0.67). For KAN, the model fit was poor, worse than a simple average of the data, i.e., slope = 0 (R^2^ < 0), indicating that it was not possible to fit the model to the KAN data. Model fit to KAN was also poor for several alternative statistical models, including Weibull, logit, logistic, and probit formulations.

**Fig. 2.**
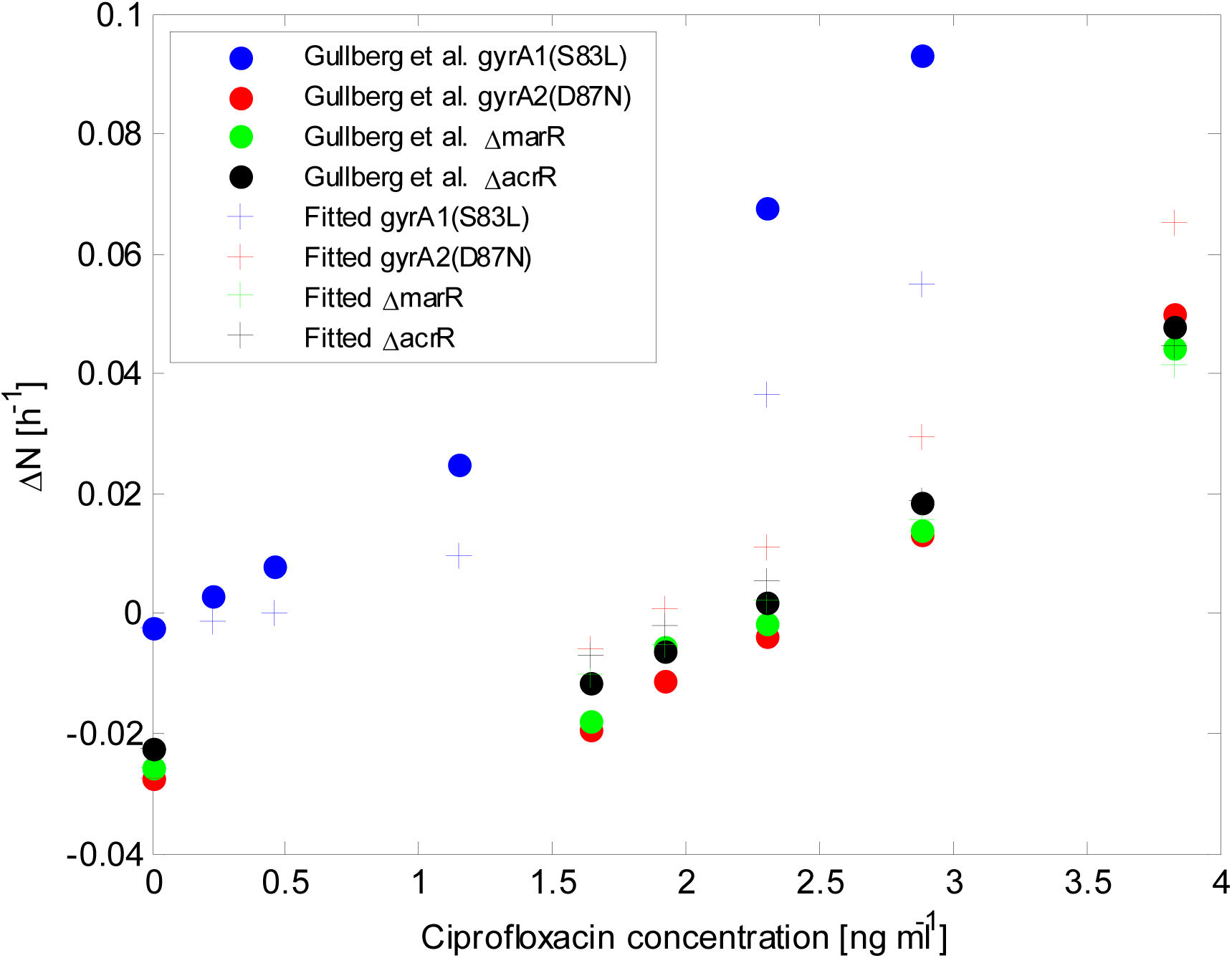
Comparison between experimentally observed (12) and model predicted difference in net growth rate between sensitive and resistant bacterial strains (ΔN, from Eq. 7) as a function of ciprofloxacin concentration in *E. coli*.

**Fig. 3.**
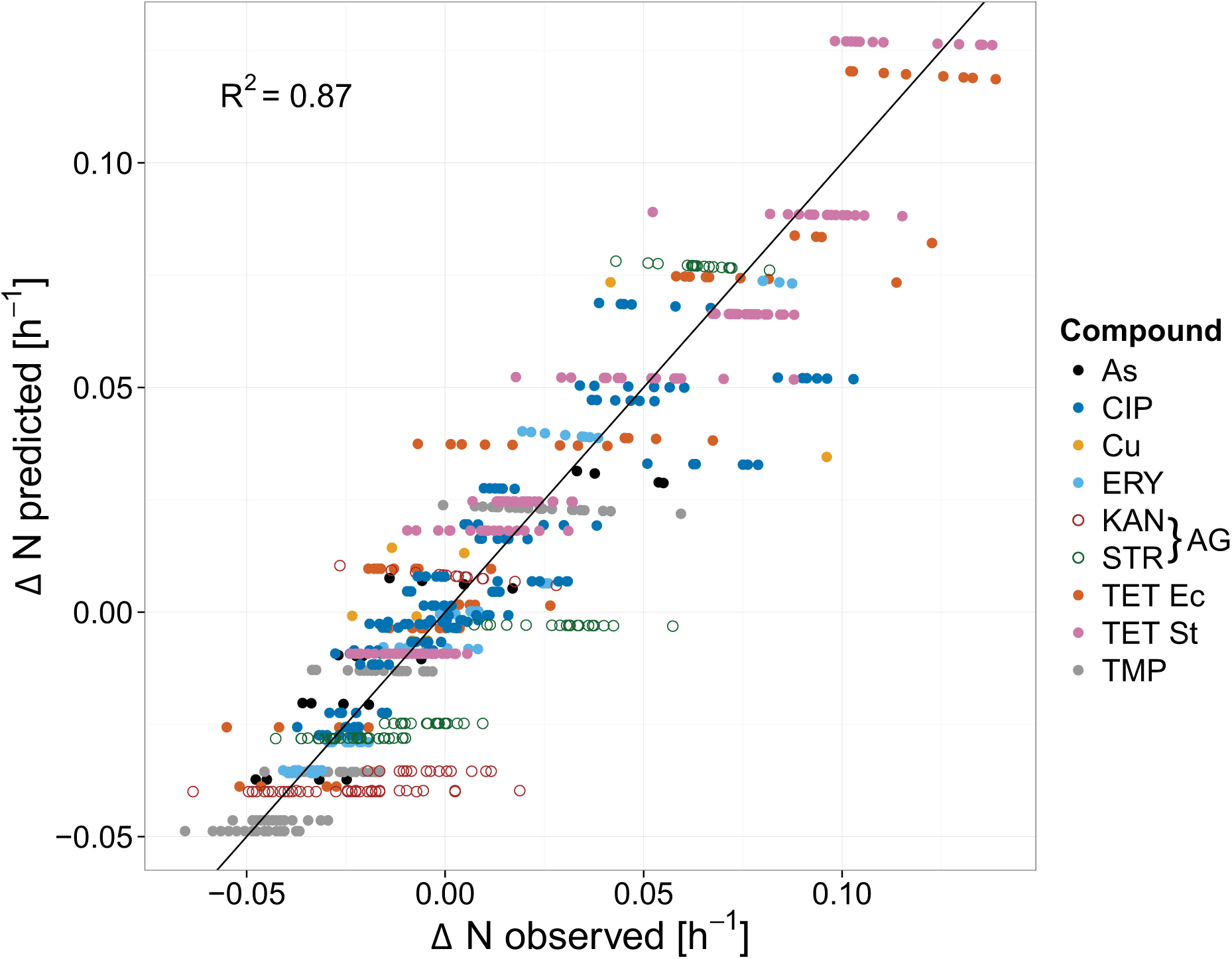
Model predicted versus ΔN observed for all experiments (12, 16). Abbreviations: As = arsenite. Cu = copper. AG = aminoglycoside antibiotic. Ec = *E. coli*. St = *S.* Typhimurium.

**Table 1.** Results of model (Eq. 7) optimization to published (12, 16) empirical growth rate differences (ΔN) between sensitive and resistant strains, based on strain-specific σ, MIC_s_, and MIC_r_. κ was fitted and raw experimental ΔN data was employed. Other model parameters: N_int,s_ = 1.8, N_min_ = −2 (19, 33, 40). MSC/MIC_s_ calculated using Eq. 10 with selection coefficients published for resistant strains (12, 16).

| Compound | Organism | Strains | n | $\kappa$ | $R^2$ | $Q^2$<sup>a</sup> | PRESS/SSY | PRESS/SSE | $MSC/MIC_s$ |
| --- | --- | --- | --- | --- | --- | --- | --- | --- | --- |
| Tetracycline<br>(TET) | *E. coli* | 3 | 60 | 1.6 | 0.89 | 0.89 | 0.09 | 1.03 | 0.014<br>0.063 |
| Trimethoprim<br>(TMP) | *E. coli* | 2 | 118 | 2.5 | 0.88 | 0.87 | 0.07 | 1.03 | 0.18 |
| Erythromycin<br>(ERY) | *E. coli* | 3 | 64 | 3.5 | 0.94 | 0.93 | 0.07 | 1.06 | 0.074<br>0.27 |
| Kanamycin<br>(KAN) | *E. coli* | 2 | 72 | 10.5 | -0.47 | -0.48 | 0.43 | 1.01 | 0.66 |
| Arsenite (As) | *E. coli* | 2 | 20 | 0.7 | 0.84 | 0.81 | 0.30 | 1.14 | 0.0064 |
| Copper sulfate<br>(Cu) | *E. coli* | 2 | 8 | 1.9 | 0.73 | 0.43 | 0.80 | 2.13 | 0.035 |
| Ciprofloxacin<br>(CIP) | *E. coli* | 5 | 144 | 2.0 | 0.78 | 0.77 | 0.31 | 1.03 | 0.024<br>0.088 |
| Streptomycin<br>(STR) | *Salmonella* | 2 | 87 | 5.0 | 0.67 | 0.66 | 0.25 | 1.02 | 0.38 |
| Tetracycline<br>(TET) | *Salmonella* | 2 | 154 | 1.2 | 0.93 | 0.93 | 0.04 | 1.02 | 0.0077 |

a. $Q^2$ = cross validated $R^2 = 1$ (PRESS/TSS)

As shown in Table 1, the fitted κ ranged widely across the nine compounds examined (0.7 to 10.5). CV results generally produced a very narrow range, with κ varying by < 0.1 within individual compounds, except KAN and Cu (supplemental material Table S2). Similarly, CV PRESS/SSY results were < 0.4 for all compounds except KAN and Cu. Values < 0.4 are considered to indicate reasonably low model prediction error (39). The PRESS/SSE were below 1.15 for all compounds except Cu; these values of PRESS/SSE close to 1 indicate limited dependence of model prediction accuracy on individual observations.

Because it was the only experiment that included four resistance genotypes, ciprofloxacin was examined more closely. Overall, fit and predictive ability were generally reasonable (R^2^ = 0.81, Q^2^ = 0.78, PRESS/SSY = 0.29, supplemental material Table S2) except for downward bias in the two highest ΔN results (Fig. 2). These were both *gyrA1* [*S83L*] versus sensitive wild-type above 2 ng/ml ciprofloxacin (19). The *gyrA1* [*S83L*] comparison had a substantially different curve shape, and removing this strain from the data greatly improved the model fit (R^2^ = 0.97, Q^2^ = 0.97, PRESS/SSY = 0.04). However the change in predicted κ was trivial (from 2.0 to 2.1, with N_min_ fixed at −2).

#### Minimum Selection Concentration

MSC/MIC_s_ was estimated (Eq. 10) based on model fitted κ, and empirical values for sc, MIC_r_, and MIC_s_. For these estimates, N_int,s_ was set at 1.8 h^-1^ and N_min_ was either fitted or set at −2 h^-1^. Model predictions corresponded well to the observed MSC/MIC_s_ (12, 16) for all experiments, with either fixed or fitted N_min_ (Fig. 4), suggesting that the model is appropriate to estimate the MSC/MIC_s_ ratio. The MSC/MIC_s_ ratio ranged across two orders of magnitude from 0.006 to 0.66 (Table 1, supplemental material Table S3).

**Fig. 4.**
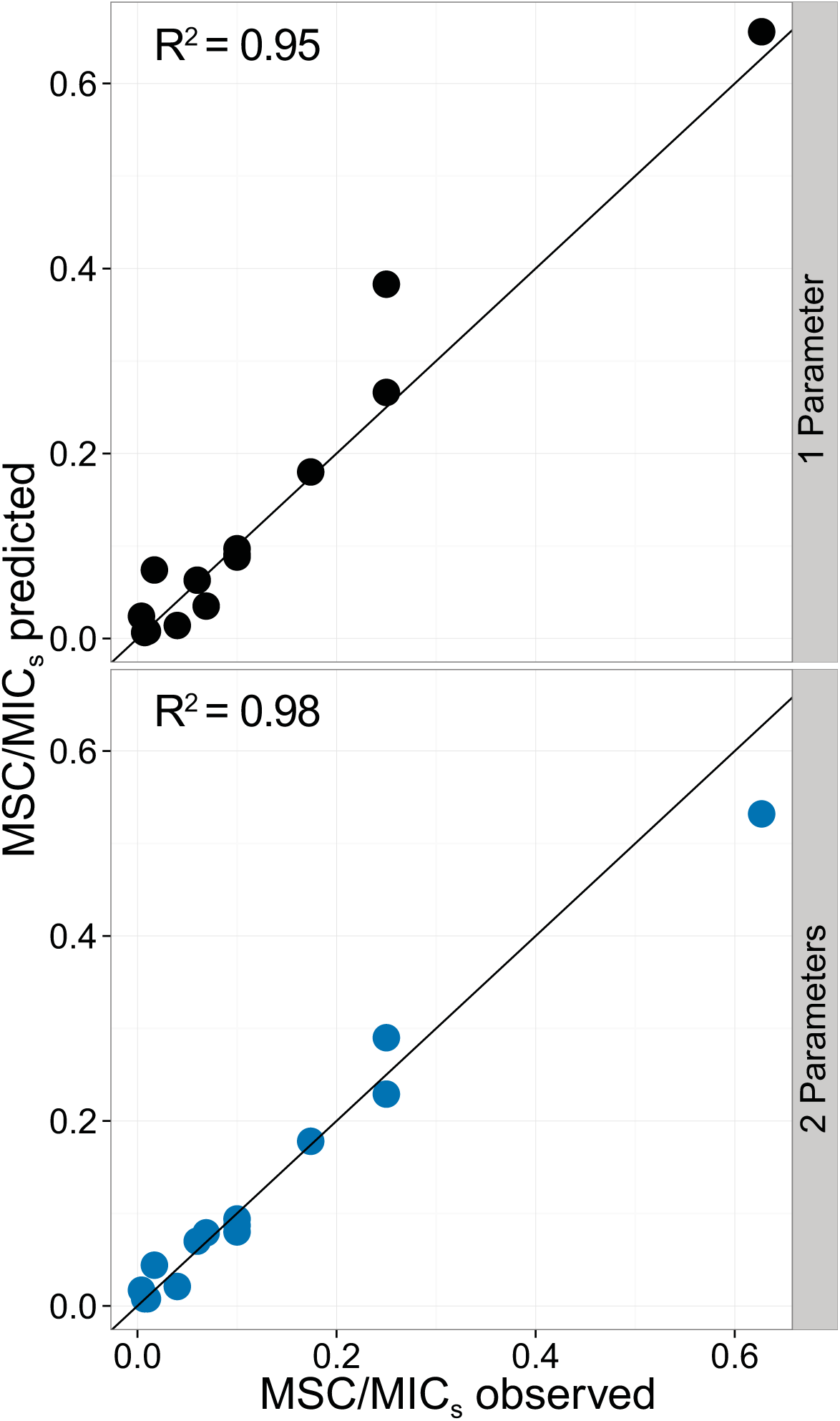
Comparison of observed (12, 16) versus model predicted (Eq. 10) MSC/MIC_s_. Symbols represent experimentally evaluated resistant strains (N = 14). Solid line (—) is 1:1 ratio.

### Sensitivity analysis

For a sensitivity analysis, behavior of Eq. 10 was examined across reasonable parameter ranges to examine sensitivity of MSC/MIC_s_ to fitness differences (sc), antibiotic resistance differences (MIC_r_/MIC_s_), maximum growth rate inhibition (N_min_), and intrinsic growth rate (N_int,s_), respectively. Eqs. 10 and 11 indicate that MSC/MIC_s_ is primarily a function of sc and κ, but is also modified by corrective terms that include N_min_, N_int,s_, N_int,r_, MIC_s_, and MIC_r_. Fig. 5 demonstrates the influences of sc and κ on MSC/MIC_s_. Specifically, increasing sc lowers the resistant strain growth rate (Fig. 5A-B), whereas increasing κ increases the curvature of the sensitive strain growth rate (Fig. 5B-D), both resulting in increased MSC/MIC_s_. As a result, modeled κ is strongly associated with model predicted MSC/MIC_s_. Thus, the Pearson correlation coefficient was very high (r = 0.94) for the κ versus MSC/MIC_s_ results from Table 1.

**Fig. 5.**
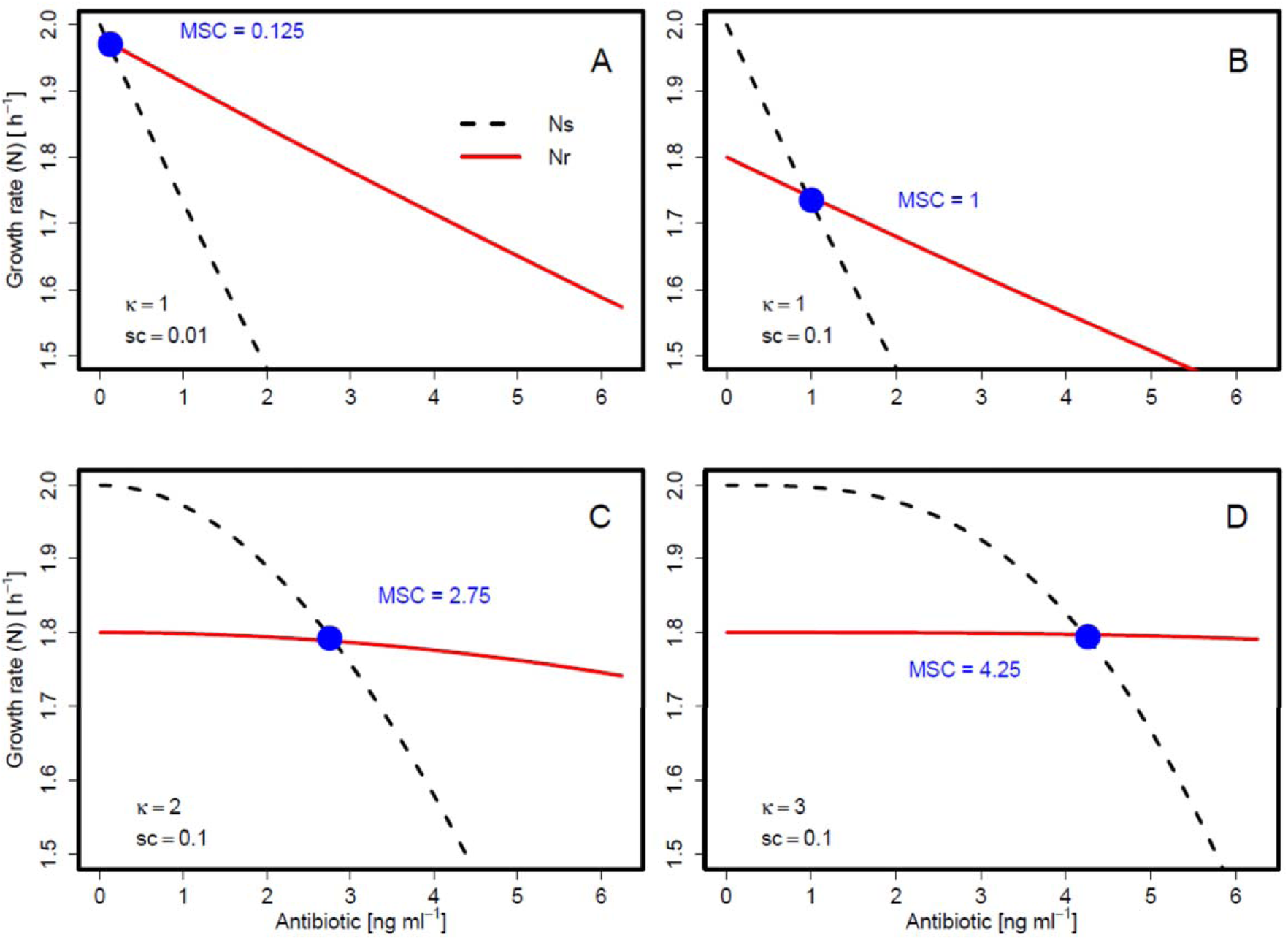
Growth rate versus antibiotic concentration for sensitive (N_s_) and resistant (N_r_) bacteria for different values of sc (0.01, 0.1) and κ (1, 2, 3). Other parameters (all scenarios): MIC_s_ = 10, MIC_r_ = 40, N_int,s_ = 2, N_min_ = −5.

Fig. 6 provides plots of the MSC/MIC_s_ ratio as the solution for Eq. 10 across different parameter values. Fig. 6A confirms the dominant and interdependent influences of sc and κ on MSC/MIC_s_ with the largest influences at or below κ values of 1. At κ = 1, MSC/MIC_s_ ≈ sc (Fig. 6A, blue dashed line). The influences of sc and κ can be combined according to Eqs. 10 and 11, which indicate that the MSC/MIC_s_ ratio is proportional to sc^1/κ^. Figs. 6B-D illustrate that MSC/MIC_s_ is proportional to sc^1/κ^ and that the slope of this relationship is modified by MIC_r_, N_int,s_, and N_min_. An increase in MIC_r_ will decrease MSC/MIC_s_, but this relationship is only sensitive when MIC_r_ approaches MIC_s_ (Fig. 6B, MIC_r_/MIC_s_ close to 1). The generally low sensitivity of MSC/MIC_s_ to the MIC values themselves corroborates the empirical finding of Gullberg et al. (12). Increasing N_int,s_ also decreases MSC/MIC_s_ but this only exhibits a minor influence in the plausible parameter range (Fig. 6C). Finally, increasing N_min_ also decreases MSC/MIC_s_, but this is only sensitive when N_min_ approaches zero (Fig. 6D). N_min_ indirectly affects MSC/MIC_s_ by influencing the MIC versus EC_50_ relationship (Eqs. S7 and S8 in supplemental material).

**Fig. 6.**
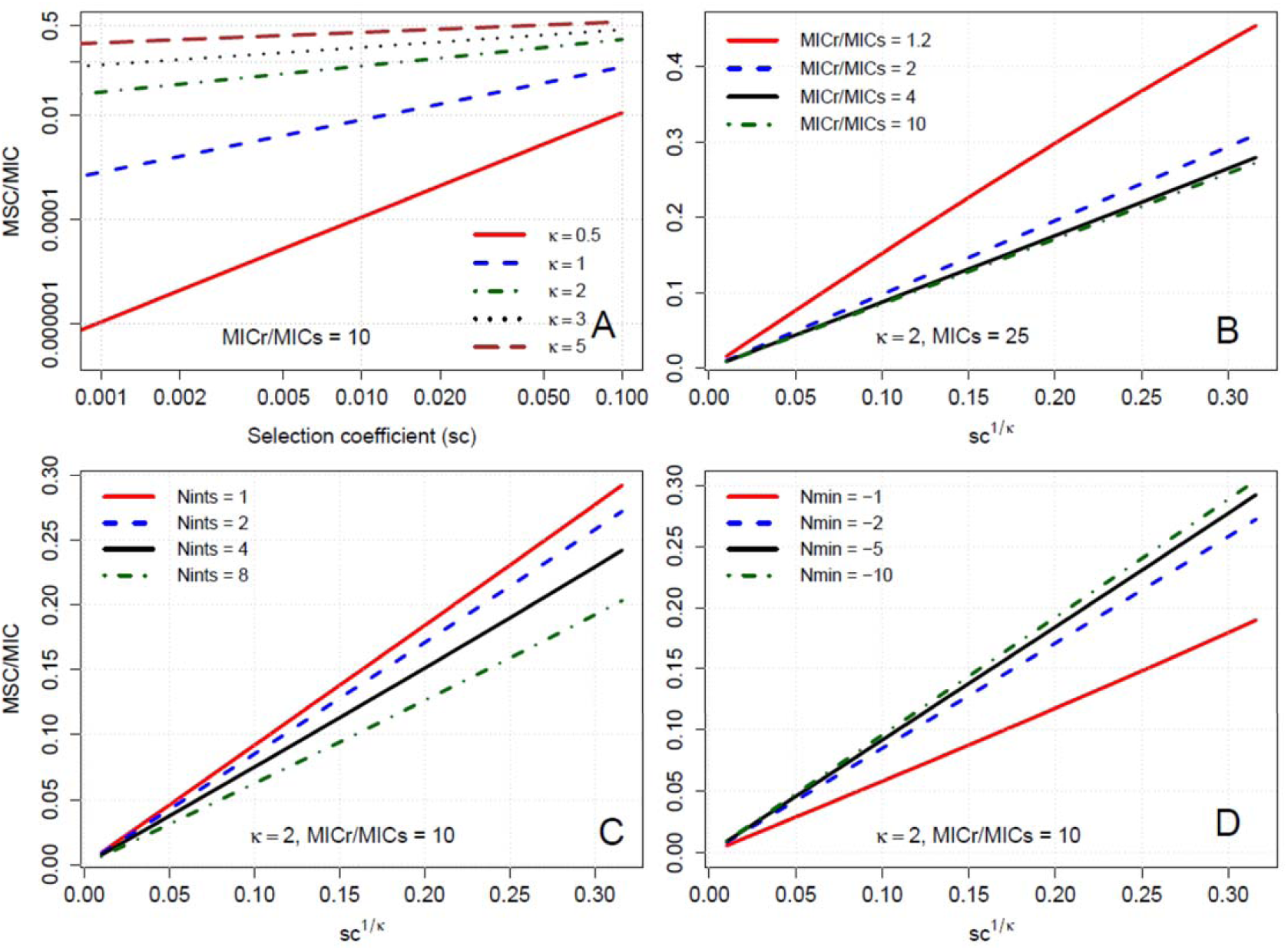
Sensitivity analysis of MSC/MIC_s_ ratio (Eq. 10) to model parameters: (A) as a function of the selection coefficients (sc) for different κ (log scale); and as a function of sc^1/^κ for different values of (B) MIC_s_/ MIC_r_, (C) N_int,s_, and (D) N_min_. Other parameters, except as noted: MIC_s_ = 25, MIC_r_ = 250, N_int,s_ = 2, N_min_ = −5. Note: panel A is log scale and panels B-D are linear scale.

Fig. 6A also illustrates the expected range of the MSC/MIC_s_ ratio across combinations of sc and κ (the most influential parameters). Over commonly observed sc ranges from 0.001 to 0.1 and κ ranges from 0.5 to 5 (12, 25, 26, 28), MSC/MIC_s_ ranged widely from 10^-6^ to 0.5. With sc = 0.01, as κ decreased from 2 to 0.5, the MSC/MIC_s_ ratio decreased from typically a factor of 0.1 down to less than a factor of 10^-4^, indicating that MSC values are very sensitive around κ = 1. Especially for low sc, slight decreases in κ may correspond to steep declines in the MSC value (Fig. 6A).

## Discussion

This study model is a simple mathematical approach to describe the factors that will drive the MSC, which is a relevant environmental threshold concentration for selection of resistant bacteria. This model helps us understand the significant issue of environmental resistance spread (1, 5, 41) by mathematically formulating the dependence of the MSC (11, 15) on the intrinsic growth rate and antibiotic versus growth dose-response. The model would enable indirect estimation of the MSC using measurements of bacterial growth parameters that are readily obtained in the laboratory and literature (κ, N_min_, and N_int_), as an alternative and possible complement to direct measurement (12, 13, 16). More importantly, the model identifies the shape of the antibiotic dose–response curve of the sensitive strain (i.e., κ) and the selection coefficient (sc) as the main parameters determining the MSC/MIC ratio. These traits, combined with literature MIC ranges (4, 8, 34), can be used to estimate environmental antibiotic concentrations at which resistance could spread.

The model consistently estimated the MSC/MIC ratio across the nine compound and taxa combinations examined, with overall R^2^ above 0.95 (Fig. 4). This finding suggests that one could estimate the MSC given: 1. the MIC; 2. intrinsic bacterial growth rate (i.e., N_int_); 3. fitness loss (either σ or sc measurements); and 4. the shape of a dose–response curve for antibiotic concentration versus bacterial growth (i.e., κ). The first three values are readily available for a range of strains, resistance mechanisms, and conditions (4, 12, 19, 25, 26, 33–36). The antibiotic dose–response curve varies across treatment conditions but is routinely obtained, allowing experimental calculation of κ (19, 21, 32, 33 provide examples).

To illustrate use of the model, Fig. 7 displays the MSC/MIC ratio from Eq. 10 across a range of selection coefficients, based on laboratory growth parameters from Regoes et al. (19) and Ankomah et al. (33). Results vary dramatically across experiments, even for the same species-antibiotic combination (Fig. 7), largely due to variations in κ. This suggests a strong impact of specific strains and growth conditions for selection, resulting in multiple orders of magnitude differences among systems, and a need to understand how the antibiotic resistance dose–response varies across antibiotic-contaminated environments (1), including water treatment systems, agricultural waste pens, and natural waters and sediments (4, 42–45).

**Fig. 7.**
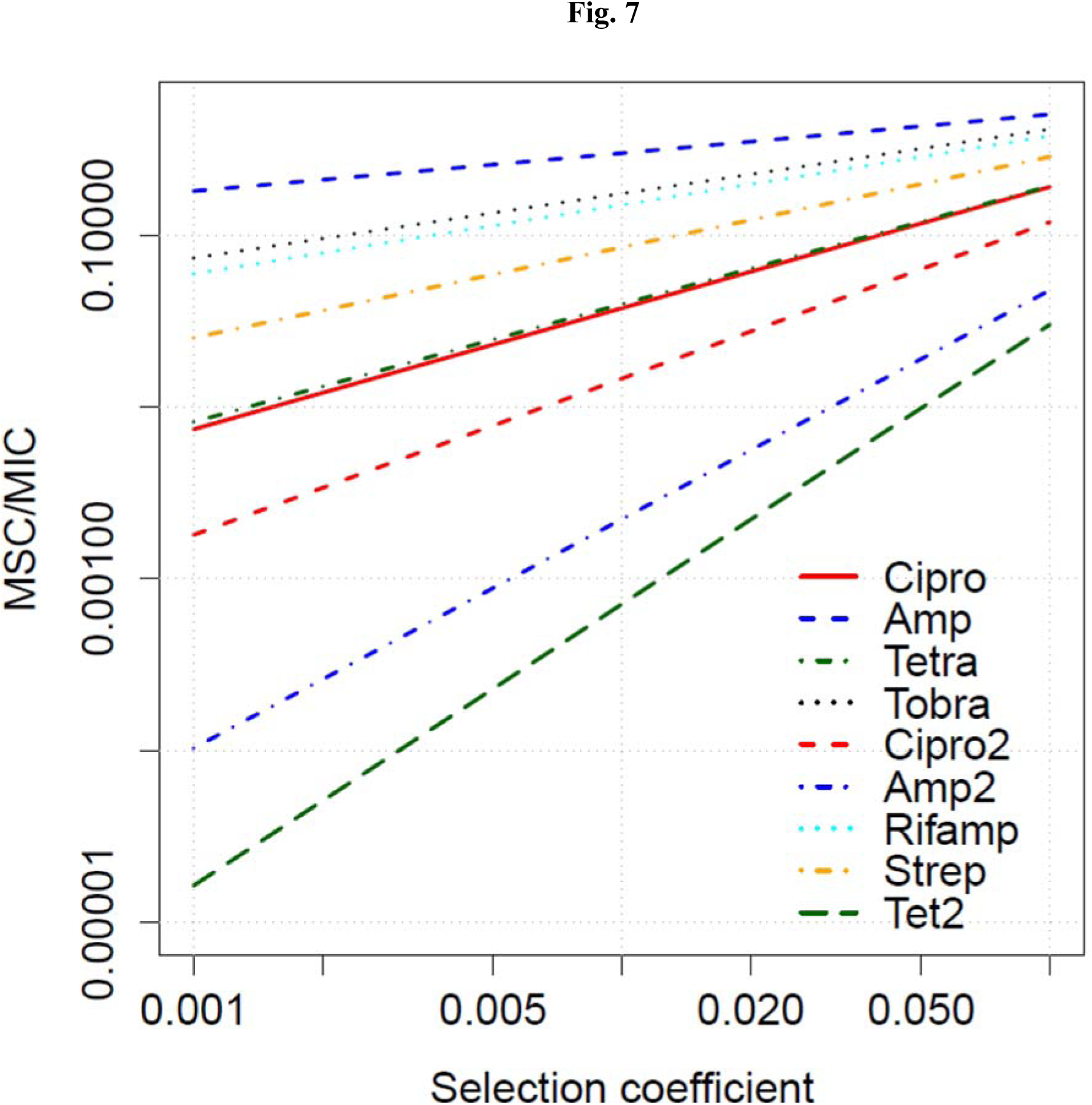
MSC/MIC as a function of the selection coefficient sc, calculated for parameters obtained in laboratory empirical studies (19, 33). Parameter data in supplemental material Table S4.

Though the model predicted the MSC/MIC well across the compounds examined, the model inconsistently predicted ΔN among compounds. ΔN was predicted least well for KAN and STR, both aminoglycosides. In these cases, the inability to fit ΔN well was due to the similarity of the study-observed MSC versus the sensitive strain MIC (i.e., high MSC/MIC_s_ ratio). This amounted to a sudden and dramatic shift from the low experimentally determined ΔN values (|ΔN| < 0.04) around the MSC versus ΔN > 1 at the MIC. This steep dose–response from high to zero growth of the sensitive strain is evident in high κ values for both STR (κ = 5) and KAN (κ = 10.5). The Hill equation and other common statistical curves could not account for the similar MSC and MIC. The high κ fitted is also inconsistent with the concentration-dependent (i.e., low κ) bactericidal activity of aminoglycoside antibiotics described elsewhere (32, 46). Instead, the similar net growth rates of susceptible versus resistant strains close to the MIC_s_ may result from adaptive resistance of the susceptible strain. Adaptive resistance for aminoglycosides has been widely observed in *Pseudomonas aeruginosa* (47, 48), including at sub-MIC exposures (49), as well as in *E. coli (22, 40, 50)*. This temporary development of phenotypic tolerance occurs due to elevated production of efflux pumps counteracting growth inhibition and killing at sublethal concentrations (49). In cases of adaptive resistance, the MSC may not be much lower than the MIC. In such cases, the MIC may be a reasonable proxy for the MSC, as is often observed clinically (7).

Experimental data are currently limited to a few species, strains, and antibiotics, possibly limiting the generalizability of the model performance evaluation. Thus, future experimental work is warranted to evaluate the ability to estimate MSC via Eqs. 10 and 11 across a range of subclinical conditions, species, strains, and antibiotics. This would include a comparison of MSC directly measured in competition experiments versus MSC derived from Eq. 10 based on measurement of the antibiotic dose–response of individual strains in isolation (Eqs. 4 and 5).

### The shape of the antibiotic dose–response at subinhibitory concentrations

By emphasizing subinhibitory antibiotic concentrations, this study extends prior findings regarding how the behavior of the Hill equation, and κ in particular, influences the dynamics of bacterial net growth (19, 32). The model predicts that an antibiotic with a lower κ for a given set of conditions (e.g., bacterial strain, media) exerts a greater selective pressure in the subinhibitory region of concentrations found in the environment, resulting in lower MSC/MIC ratios. With κ ≈ 1, there is an approximately linear decrease in growth from the intrinsic rate with no antibiotic to zero growth when the antibiotic concentration is equal to the MIC. As a result, the intersection between the curves for the wild-type versus resistant strain can occur at a low antibiotic concentration, and the MSC is approximately equal to the MIC of the wild-type multiplied by the selection coefficient. This leads to a low MSC for low selection coefficients.

For higher κ conditions, the MSC is closer to the MIC. Thus, high κ, in addition to increasing efficacy above the MIC (19), also reduces the hazard of selection for resistance at concentrations below the MIC. Simulated and empirical dose–response measurements in the subinhibitory region are especially needed to evaluate the extent to which that ‘pre-selection’ of resistant strains may occur at MSC levels below the MIC of the sensitive strain, in both clinical and environmental settings.

### Implications for resistance development hazard

Environmental-hazard and -risk assessments would benefit from determining how ambient environmental concentrations in different media compare to the MSC (1). Based on a species sensitivity distribution compared to EUCAST-published MIC results, Tello et al. found that selective pressure for resistant bacterial communities would be high in swine feces lagoon sediment but low in surface water, ground water, raw sewage, and sewage treatment plant effluent (4). As an example of the implications of the MSC threshold (versus the MIC), we reinterpret the model of Tello et al. (4) to estimate hazard of selection for resistant bacteria. We employ a model correction factor, assuming a 100-fold lower species sensitivity distribution, to convert the study reported MIC_50_ (Fig. 4 in Ref. 4) to an MSC_50_ by adjusting the reported log-logistic model location (α) parameter by −2. The 100-fold reduction follows our model results and the empirical data of Gullberg et al. (12, 16), both of which indicate MSC/MIC ratios may exhibit values below 0.01. Comparing the adjusted model to the field data reported for ciprofloxacin (Table 2 in Ref. 4), the MSC_50_ model predicted a greater than 25% potentially affected fraction of bacterial taxa in at least one sample for all media reported (surface water, river sediment, raw sewage, and treatment plant effluent). For erythromycin and tetracycline, the MSC_50_ model predicted 65% and 88% potentially affected fraction in river sediment (vs. 2% and 1.6% for the MIC_50_) (4). Tello et al. used data from systems impacted by human and agricultural development (42, 43), and our 100-fold MSC:MIC correction is more conservative than a 10-fold reduction employed in PNECs developed by Bengtsson-Palme and Larsson (8), thus indicating worst-case conditions. Nevertheless, Bengtsson-Palme and Larsson also reported treatment plant effluent concentrations above PNECs (8) in 28% of cases. These results in combination indicate likely selection of resistant strains given antibiotic exposure in a wide variety of human-impacted aquatic settings.

### Model scope, limitations, and future directions

The parsimonious analytical solution we developed addresses vertical gene transfer of antibiotic resistance in a well-mixed environment as a function of fitness loss, competition, and antibiotic concentration. There are many aspects of resistance dissemination that fall outside the scope of this simple exercise, including horizontal gene transfer, interactions among multiple strains, spatial arrangement of individual colonies, and heterogeneity in antibiotic exposure due to biofilms and other mechanisms (2, 8, 11, 20, 51). Additionally, the model operates on and describes the long-term competition dynamics between bacterial strains, rather than stochastic and dynamic changes in net growth and competition over time. Thus, the derivation assumes that the parameters governing growth (e.g., R_int_, D_int_, D_ab_) will reach relatively stable values when one strain outcompetes another strain. This simplified model does not include parameters for the inoculum effect, biphasic killing, delay functions, drug concentration changes, drug-insusceptible persister cells, or adaptive resistance, all of which may occur in experimental settings. More sophisticated pharmacokinetic-pharmacodynamic models, which incorporate these processes are needed to characterize the bacterial time-kill curve and optimal dosing regimens (21–23). However, such models do not lend themselves to an analytical solution similar to what we have provided. Investigation of varying initial ratios of resistant versus susceptible bacteria indicate no effect on selection coefficient, suggesting a limited importance of initial conditions, such as inoculum effect (12, 16). Nevertheless, theoretical and experimental investigation of how short-term growth and killing and other dynamic processes would impact the MSC/MIC ratio is warranted in future studies, as is comparison of alternative models.

The primary benefit of the present model is in illustrating the MSC paradigm and the key drivers of selection in simplified systems. As such, this paper adds to the growing scientific understanding on how to interpret laboratory data on the MIC and other parameters for predicting the emergence of resistance at subinhibitory environmental concentrations. It highlights the value of characterizing the antibiotic dose-response (i.e., the Hill Coefficient κ), particularly at antibiotic concentrations below the MIC. Ultimately, this quantification of resistance selection must be integrated into a risk assessment framework that also considers environmental antibiotic contamination, human exposure to and colonization by resistant bacteria, and the association between colonization and infection (1). Such a framework can help quantify the global hazard posed by antimicrobial agents.

## Acknowledgements

This research has been partly carried out as part of the SUBMERGE program at University of Michigan, with support of the Graham Environmental Sustainability Institute at University of Michigan. BG was supported by a US EPA STAR Fellowship (Environmental Protection Agency FP917287) and a SAGE-IGERT traineeship (National Science Foundation Award Number 1144885). We thank Scott Reed for performing valuable analyses, and Peter Adriaens, Cedric Wannaz (University of Michigan) and Lee Riley (UC Berkeley) for helpful comments and input. Review by Lee Riley, John Balmes, and three anonymous reviewers greatly improved a previous version of the manuscript.

